# Identification of Plitidepsin as Potent Inhibitor of SARS-CoV-2-Induced Cytopathic Effect after a Drug Repurposing Screen

**DOI:** 10.1101/2020.04.23.055756

**Authors:** Jordi Rodon, Jordana Muñoz-Basagoiti, Daniel Perez-Zsolt, Marc Noguera-Julian, Roger Paredes, Lourdes Mateu, Carles Quiñones, Itziar Erkizia, Ignacio Blanco, Alfonso Valencia, Víctor Guallar, Jorge Carrillo, Julià Blanco, Joaquim Segalés, Bonaventura Clotet, Júlia Vergara-Alert, Nuria Izquierdo-Useros

## Abstract

There is an urgent need to identify therapeutics for the treatment of Coronavirus diseases 2019 (COVID-19). Although different antivirals are given for the clinical management of SARS-CoV-2 infection, their efficacy is still under evaluation. Here, we have screened existing drugs approved for human use in a variety of diseases, to compare how they counteract SARS-CoV-2-induced cytopathic effect and viral replication *in vitro.* Among the potential 72 antivirals tested herein that were previously proposed to inhibit SARS-CoV-2 infection, only 18% had an IC_50_ below 25 μM or 10^2^ IU/mL. These included plitidepsin, novel cathepsin inhibitors, nelfinavir mesylate hydrate, interferon 2-alpha, interferon-gamma, fenofibrate, camostat along the well-known remdesivir and chloroquine derivatives. Plitidepsin was the only clinically approved drug displaying nanomolar efficacy. Four of these families, including novel cathepsin inhibitors, blocked viral entry in a cell-type specific manner. Since the most effective antivirals usually combine therapies that tackle the virus at different steps of infection, we also assessed several drug combinations. Although no particular synergy was found, inhibitory combinations did not reduce their antiviral activity. Thus, these combinations could decrease the potential emergence of resistant viruses. Antivirals prioritized herein identify novel compounds and their mode of action, while independently replicating the activity of a reduced proportion of drugs which are mostly approved for clinical use. Combinations of these drugs should be tested in animal models to inform the design of fast track clinical trials.

## INTRODUCTION

A novel betacoronavirus, the severe acute respiratory syndrome coronavirus 2 (SARS-CoV-2), is causing a respiratory disease pandemic that began in Wuhan, China, in November 2019, and has now spread across the world^1^. To date, remdesivir is the only approved antiviral drug for the specific treatment of this coronavirus infectious disease 2019 or COVID-19^2,3^. However, several drugs are being used in the frontline of clinical management of SARS-CoV-2-infected individuals in hospitals all around the world, to try to avoid the development of the COVID-19 associated pneumonia, which can be fatal. By the end of December 2020, almost 1.7 million people had died from COVID-19, and over 75 million cases have been reported (https://covid19.who.int).

Although different drug regimens are being applied to hospitalized patients, no clinical study has evidenced their efficacy yet. Under this scenario, initiatives launched by the World Health Organization (WHO), such as the SOLIDARITY study that has compared remdesivir, hydroxychloroquine, ritonavir/lopinavir and ritonavir/lopinavir plus β-interferon regimes, have been of critical importance to prioritize the use of the most active compounds^4^ Unfortunately, although remdesivir has proven efficacy in randomized controlled trials^2,3^, a recent update of the WHO clinical trial has failed to detect any effect on overall mortality, initiation of ventilation and duration of hospital stay with any of the antivirals tested^5^. Thus, there is an urgent need to identify novel therapeutic approaches for individuals with COVID-19 developing severe disease and fatal outcomes.

In this report we present a prioritized list of effective compounds with proven antiviral efficacy *in vitro* to halt SARS-CoV-2 replication. Compounds were analyzed depending on their expected mechanism of action, to identify candidates tackling diverse steps of the viral life cycle. SARS-CoV-2 entry requires viral binding and spike protein activation via interaction with the cellular receptor ACE2 and the cellular protease TMPRSS2^6^, a mechanism favored by viral internalization via endocytosis. Interference with either of these processes has proven to decrease SARS-CoV-2 infectivity^6,7^, and therefore, inhibitors targeting viral entry may prove valuable. In addition, SARS-CoV-2 enters into the cells via endocytosis and accumulates in endosomes where cellular cathepsins can also prime the spike protein and favor viral fusion upon cleavage^6,8,9^, providing additional targets for antiviral activity. Once SARS-CoV-2 fuses with cellular membranes, it triggers viral RNA release into the cytoplasm, where polyproteins are translated and cleaved by proteases^10^. This leads to the formation of an RNA replicase-transcriptase complex driving the production of several negative-stranded RNA via both replication and transcription^10^. Numerous negative-stranded RNAs transcribe into messenger RNA genomes, allowing for the translation of viral nucleoproteins, which assemble in viral capsids at the cytoplasm^10^. These capsids then bud into the lumen of endoplasmic reticulum (ER)-Golgi compartments, where viruses are finally released into the extracellular space by exocytosis. Potentially, any of these viral cycle steps could be targeted with antivirals, so we have thus searched for these compounds as well.

Finally, as the most effective antiviral treatments are usually based on combined therapies that tackle distinct steps of the viral life cycle, we also tested the active compounds in combination. These combinations may be critical to abrogate the potential emergence of resistant viruses and to increase antiviral activity, enhancing the chances to improve clinical outcome.

## MATERIAL & METHODS

### Ethics statement

The institutional review board on biomedical research from Hospital Germans Trias i Pujol (HUGTiP) approved this study. The individual who provided the sample to isolate virus gave a written informed consent to participate.

### Cell Cultures

Vero E6 cells (ATCC CRL-1586) were cultured in Dulbecco’s modified Eagle medium, (DMEM; Lonza) supplemented with 5% fetal calf serum (FCS; EuroClone), 100 U/mL penicillin, 100 μg/mL streptomycin, and 2 mM glutamine (all ThermoFisher Scientific). HEK-293T (ATCC repository) were maintained in DMEM with 10% fetal bovine serum, 100 IU/mL penicillin and 100 μg/mL streptomycin (all from Invitrogen). HEK-293T overexpressing the human ACE2 were kindly provided by Integral Molecular Company and maintained in DMEM (Invitrogen) with 10% fetal bovine serum, 100 IU/mL penicillin and 100 μg/mL streptomycin, and 1 μg/mL of puromycin (all from Invitrogen). TMPRSS2 human plasmid (Origene) was transfected using X-tremeGENE HP Transfection Reagent (Merck) on HEK-293T overexpressing the human ACE2 and maintained in the previously described media containing 1 mg/ml of geneticin (Invitrogen) to obtain TMPRSS2/ACE2 HEK-293T cells.

### Virus isolation, titration and sequencing

SARS-CoV-2 was isolated from a nasopharyngeal swab collected from an 89-year-old male patient giving informed consent and treated with Betaferon and hydroxychloroquine for 2 days before sample collection. The swab was collected in 3 mL medium (Deltaswab VICUM) to reduce viscosity and stored at −80°C until use. Vero E6 cells were cultured on a cell culture flask (25 cm^2^) at 1.5 x 10^6^ cells overnight prior to inoculation with 1 mL of the processed sample, for 1 h at 37°C and 5% CO_2_. Afterwards, 4 mL of 2% FCS-supplemented DMEM were supplied and cells were incubated for 48 h. Supernatant was harvested, centrifuged at 200 x g for 10 min to remove cell debris and stored at −80°C. Cells were assessed daily for cytopathic effect and the supernatant was subjected to viral RNA extraction and specific RT-qPCR using the SARS-CoV-2 UpE, RdRp and N assays^11^. The virus was propagated for two passages and a virus stock was prepared collecting the supernatant from Vero E6.

Viral RNA was extracted directly from the virus stock using the Indimag Pathogen kit (Indical Biosciences) and transcribed to cDNA using the PrimeScript™ RT reagent Kit (Takara) using oligo-dT and random hexamers, according to the manufacturer’s protocol. DNA library preparation was performed using SWIFT amplicon SARS-CoV-2 panel (Swift Biosciences). Sequencing ready libraries where then loaded onto Illumina MiSeq platform and a 300bp paired-end sequencing kit. Sequence reads were quality filtered and adapter primer sequences were trimmed using trimmomatic. Amplification primer sequences were removed using cutadapt^12^ Sequencing reads were then mapped against coronavirus reference (NC_045512.2) using bowtie2 tool^13^. Consensus genomic sequence was called from the resulting alignment at a 18×1800×879 average coverage using samtools^14^ Genomic sequence was deposited at GISAID repository (http://gisaid.org) with accession ID EPI_ISL_510689.

### Pseudovirus production

HIV-1 reporter pseudoviruses expressing SARS-CoV-2 Spike protein and luciferase were generated using two plasmids. pNL4-3.Luc.R-.E-was obtained from the NIH AIDS repository. SARS-CoV-2.SctΔ19 was generated (Geneart) from the full protein sequence of SARS-CoV-2 spike with a deletion of the last 19 amino acids in C-terminal, human-codon optimized and inserted into pcDNA3.4-TOPO^15^. Spike plasmid was transfected with X-tremeGENE HP Transfection Reagent (Merck) into HEK-293T cells, and 24 hours post-transfection, cells were transfected with pNL4-3.Luc.R-.E-. Supernatants were harvested 48 hours later, filtered with 0.45 μM (Millex Millipore) and stored at −80°C until use. Control pseudoviruses were obtained by replacing the spike plasmid by a VSV-G plasmid (kindly provided by Dr. Andrea Cimarelli). The p24^gag^ content of all viruses was quantified using an ELISA (Perkin Elmer) and viruses were titrated in HEK-293T overexpressing the human ACE2.

### Antivirals & compounds

The complete list of compounds used for this study and vendors are shown in Supplementary Tables 1 to 5. Drugs were used at concentrations ranging from 100 μM to 0.0512 nM at 5-fold serial dilutions. NPOs were used at concentrations ranging from 10 μM to 0.00512 nM at 5-fold serial dilutions. Plitidepsin was also assayed at concentrations ranging from 10 μM to 0.5 nM at 3-fold dilutions. Interferons were assayed at concentrations ranging from 10^4^ to 0.0005 IU/ml at 5-fold serial dilutions. When two drugs were combined, each one was added at a 1:1 molar ratio at concentrations ranging from 100 μM to 0.0512 nM at 5-fold serial dilutions. In combination with other drugs, plitidepsin was also assayed at concentrations ranging from 10 μM to 0.5 nM at 3-fold dilutions.

### Antiviral activity

Increasing concentrations of antiviral compounds were added to Vero E6 cells and immediately after, we added 10^1.8^ TCID50/mL of SARS-CoV-2, a concentration that achieves a 50% of cytopathic effect. Untreated non-infected cells and untreated virus-infected cells were used as negative and positive controls of infection, respectively. To detect any drug-associated cytotoxic effect, Vero E6 cells were equally cultured in the presence of increasing drug concentrations, but in the absence of virus. Cytopathic or cytotoxic effects of the virus or drugs were measured 3 days after infection, using the CellTiter-Glo luminescent cell viability assay (Promega). Luminescence was measured in a Fluoroskan Ascent FL luminometer (ThermoFisher Scientific).

### IC_50_ calculation and statistical analysis

Response curves of compounds or their mixes were adjusted to a non-linear fit regression model, calculated with a four-parameter logistic curve with variable slope. Cells not exposed to the virus were used as negative controls of infection, and were set as 100% of viability to normalize data and calculate the percentage of cytopathic effect. Statistical differences from 100% were assessed with a one sample t test. All analyses and figures were generated with the GraphPad Prism v8.0b Software.

### In silico drug modeling

We performed Glide docking using an in-house library of all approved drug molecules on the 3CL protease of SARS-CoV-2. For this, two different receptors were used, the 6LU7 pdb structure, after removing the covalently bound inhibitor, and a combination of two crystals from the Diamond collection (https://www.diamond.ac.uk/covid-19). Receptors were prepared with the Schrodinger’s protein wizard and Glide SP docking was performed with two different hydrogen bond constraints: Glu16 and His 163 (with epsilon protonation); we enforced single constraints and also attempted the combination of both. The best 11 molecules, based on Glides’s docking score were selected. Top docking scores, however, did not exceed −9, indicating poor potential binding.

### Pseudovirus assay

HEK-293T overexpressing the human ACE2 and TMPRSS2 were used to test antivirals at the concentrations found to be effective for SARS-CoV-2 without toxicity, which were the following: 5 μM for niclosamide; 10 μM for chloroquine, chlorpromazine, ciclesonide, MDL 28170 and fenofibrate; 20 μM for hydroxychloroquine, CA-074-Me and arbidol HCl; 25 μM for E-64d; 50 μM for Baricitinib; 100 μM for Amantadine, NB-DNJ, 3’ sialyl-lactose Na salt, Tofacitinib, and Camostat mesylate; 1000 μM for methyl-β-cyclodextrin, and 12,5 mg/ml for AAT. A constant pseudoviral titer was used to pulse cells in the presence of the drugs. At 48h post-inoculation, cells were lysed with the Glo Luciferase system (Promega). Luminescence was measured with an EnSight Multimode Plate Reader (Perkin Elmer).

### SARS-CoV-2 detection in the supernatant of infected cells

Viral accumulation in the supernatant of Vero E6 cells infected as described previously in the presence of increasing concentrations of the indicated antiviral compounds was measured at day 3 post-infection. The amount of SARS-CoV-2 nucleoprotein released to the supernatant was measured with an ELISA (SinoBiologicals), according to the manufacturer’s protocol.

## RESULTS

We have tested the antiviral activity of different clinically available compounds and their combinations by assessing their ability to inhibit viral induced cytopathic effect *in vitro.* Our strategy was circumscribed mostly to compounds approved for clinical use, since they are ideal candidates for entering into fast track clinical trials. Drug selection criteria first focused on compounds already being tested in clinical trials, along with well-known human immunodeficiency virus-1 (HIV-1) and hepatitis C virus (HCV) inhibitors, as well as other compounds suggested to have potential activity against SARS-CoV-2 in molecular docking analysis or *in vitro* assays.

We first assessed the activity of 16 compounds with hypothetical capacity to inhibit viral entry, and then we focused on 22 drugs thought to block viral replication upon SARS-CoV-2 fusion. Molecular docking studies provided an additional 11 candidates, which were predicted to inhibit the SARS-CoV-2 main protease. Finally, 23 compounds with unknown mechanism of action were also assessed. By these means, we have compared 72 drugs and 28 of their combinations for their capacity to counteract SARS-CoV-2-induced cytopathic effect *in vitro*.

### 1. Antiviral activity of compounds that potentially inhibit viral entry

We first tested compounds that could have an effect before viral entry by impairing virus-cell fusion (**Supp. Table 1**). We confirmed the inhibitory effect of hydroxychloroquine against SARS-CoV-2-induced cellular cytotoxicity on Vero E6 cells^16^. As shown in **Fig. 1A**, this drug was able to inhibit viral-induced cytopathic effects at concentrations where no cytotoxic effects of the drug were observed, as previously reported^16,17^. Since hydroxychloroquine was first administered in combination with the antibiotic azithromycin^22^, which induces antiviral responses in bronchial epithelial cells^23^, we tested this antibiotic alone and in combination (**Fig. 1A**), but found a similar activity to that of the chloroquine derivative alone (**Fig. 1A**). Indeed, this was also the case when we tested hydroxychloroquine in combination with different HIV-1 protease inhibitors and other relevant compounds currently being tested in clinical trials (**Supp. Table 2**).

**Figure 1.**
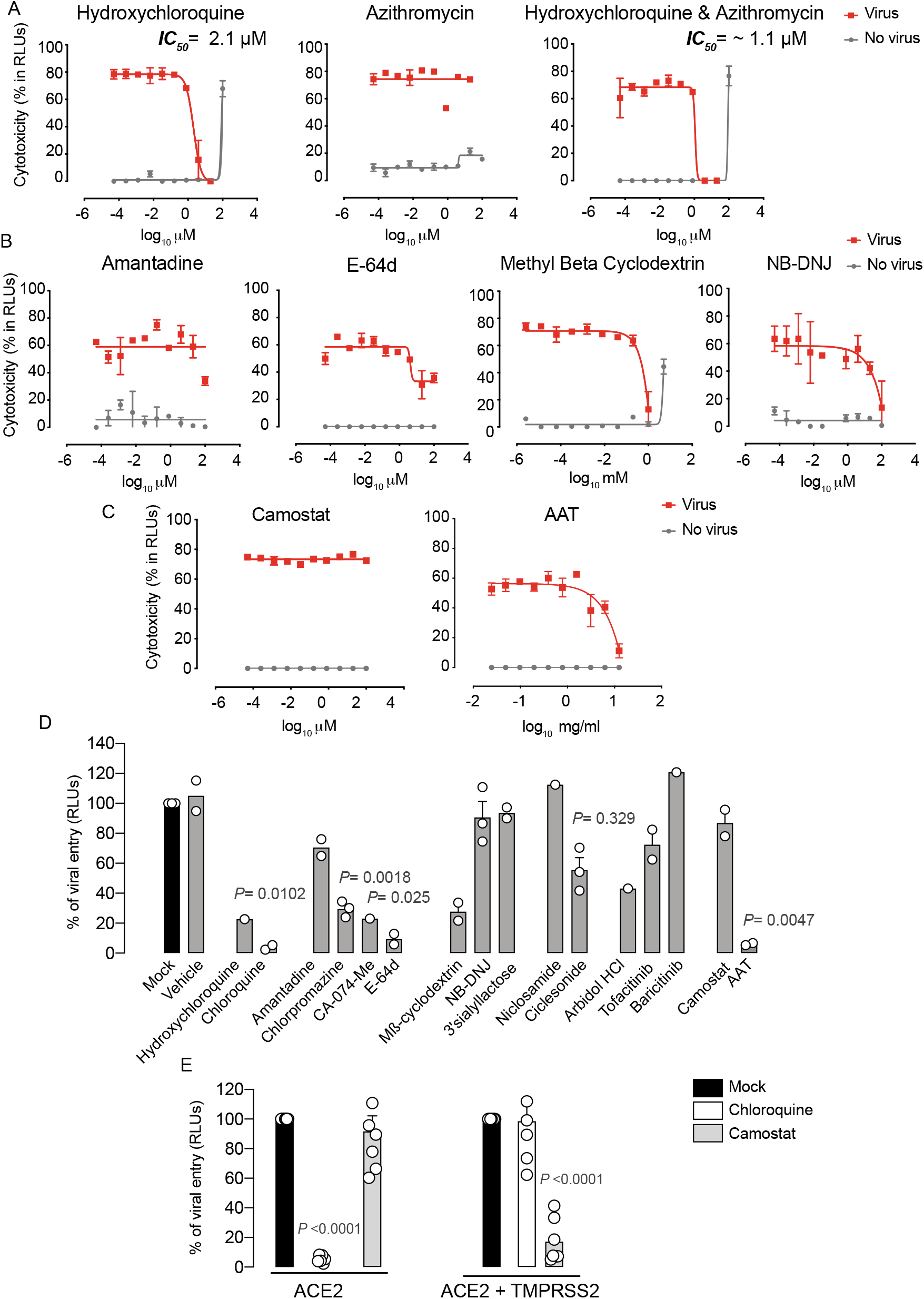
Antiviral activity of entry inhibitors against SARS-CoV-2. **A.** Antiviral activity of hydroxychloroquine and azithromycin. Cytopathic effect on Vero E6 cells exposed to a fixed concentration of SARS-CoV-2 in the presence of increasing concentrations of hydroxychloroquine, azithromycin, and their combination. Drugs were used at a concentration ranging from 0.0512 nM to 100 μM. When combined, each drug was added at the same concentration. Non-linear fit to a variable response curve from one representative experiment with two replicates is shown (red lines), excluding data from drug concentrations with associated toxicity. The particular IC_50_ value of this graph is indicated. Cytotoxic effect on Vero E6 cells exposed to increasing concentrations of drugs in the absence of virus is also shown (grey lines). **B.** Cytopathic effect on Vero E6 cells exposed to a fixed concentration of SARS-CoV-2 in the presence of increasing concentrations of amantadine, a clathrin-mediated endocytosis inhibitor, E-64d, a pan-cathepsin inhibitor acting downstream once viruses are internalized in endosomes, NB-DNJ, an inhibitor of ganglioside biosynthesis and methyl-β-cyclodextrin, a cholesterol-depleting agent. All drugs were used at a concentration ranging from 0.0512 nM to 100 μM aside from methyl-β-cyclodextrin, which was used 10 times more concentrated. Non-linear fit to a variable response curve from one experiment with two replicates is shown (red lines). Cytotoxic effect on Vero E6 cells exposed to increasing concentrations of drugs in the absence of virus is also shown (grey lines). **C.** Cytopathic effect on Vero E6 cells exposed to a fixed concentration of SARS-CoV-2 in the presence of increasing concentrations of camostat, a TMPRSS2 inhibitor, and ATT, an alpha-1 antyitrypsin, a broad cellular protease inhibitor, as described in **A. D.** Effect of entry inhibitors on luciferase expression of reporter lentiviruses pseudotyped with SARS-CoV-2 Spike in ACE2 expressing HEK-293T cells. Values are normalized to luciferase expression by mock-treated cells set at 100%. Mean and s.e.m. from two experiments with one to three replicates. Cells were exposed to fixed amounts of SARS-CoV-2 Spike lentiviruses in the presence of a nontoxic constant concentration of the drugs tested on Vero E6. Significant statistical deviations from 100% were assessed with a one sample t test. **E.** Comparison of entry inhibitors blocking viral endocytosis, such as chloroquine, with inhibitors blocking serine protease TMPRSS2 expressed on the cellular membrane, such as camostat, on different cell lines. ACE2 expressing HEK-293T cells transfected or not with TMPRSS2 were exposed to SARS-CoV-2 Spike lentiviruses as described in **B**. Values are normalized to luciferase expression by mock-treated cells set at 100%. Mean and s.e.m. from at least two representative experiments with two replicates. Statistical deviations from 100% were assessed with a one sample t test.

Additional Food and Drug Administration (FDA)-approved compounds previously used to abrogate viral entry via clathrin-mediated endocytosis such as amantadine or chlorpromazine were also tested in this SARS-CoV-2-induced cytotoxicity assay (**Supp. Table 1**). Yet, we did not find any prominent effect of these agents; only a partial inhibition at 100 μM for amantadine (**Fig. 1B**). The broad cathepsin B/L inhibitor E64-d also showed partial inhibitory activity (**Fig. 1B**). E64-d exerts activity against viruses cleaved by cellular cathepsins upon endosomal internalization, as previously described using pseudotyped SARS-CoV-2^6^. While these results could not be confirmed using the specific cathepsin B inhibitor CA-074-Me due to drug-associated toxicity (**Supp. Table 1**), none of these cathepsin inhibitors is approved for clinical use. It has also been suggested that hydroxychloroquine could block SARS-CoV-2 spike interaction with GM1 gangliosides^28^. GM1 gangliosides are enriched in cholesterol domains of the plasma membrane and have been previously shown to bind to SARS-CoV spike protein^29^ This mode of viral interaction is aligned with the capacity of methyl-beta cyclodextrin, which depletes cholesterol from the plasma membrane to abrogate SARS-CoV-2 induced cytopathic effect (**Fig. 1B**), as previously reported for SARS-CoV^29^ Removal of cholesterol redirected ACE2 receptor to other domains, but did not alter the expression of the viral receptor^29^ Moreover, NB-DNJ, an inhibitor of ganglioside biosynthesis pathway, also decreased SARS-CoV-2 cytopathic effect (**Fig. 1B**). These results highlight the possible role of gangliosides in viral binding, although in soluble competition the polar head group of GM3 ganglioside (3’ Sialyllactose) was not able to reduce viral-induced cytopathic effect (**Supp. Table 1**). Of note, previously reported antiviral agents such as the autophagy BECN1-stabilizing compounds niclosamide or ciclesonide^30,31^, the membrane lipid intercalating compound arbidol^32^ and the two JAK inhibitors baricitinib and tofacitinib^34,35^ failed to protected Vero E6 cells from SARS-CoV-2 induced cytopathic effect (**Supp. Table 1**). We next tested camostat, a serine protease inhibitor with capacity to abrogate SARS-CoV-2 Spike priming on the plasma membrane of human pulmonary cells and avoid viral fusion^6^. Camostat showed no antiviral effect on Vero E6 cells (**Fig. 1C**), what indicates that the alternative viral endocytic route is the most prominent entry route in this renal cell type. A broader cellular protease inhibitor such as alpha-1 antitrypsin (AAT), used to treat severe AAT human deficiency and currently under clinical study for COVID-19 due to its anti-inflammatory potential, has already shown capacity to limit SARS-CoV-2 entry in cells expressing TMPRSS2^36,37^ When we tested AAT antiviral efficacy on Vero E6, AAT exerted an effect but required high concentrations that will most likely rely on the activity of these proteases in the endosomal route (**Fig. 1C**).

To confirm that identified compounds listed in **Supp. Table 1** specifically inhibit the viral entry step, we employed a luciferase-based assay using pseudotyped lentivirus expressing the spike protein of SARS-CoV-2, which allows to detect viral fusion on HEK-293T cells transfected with ACE2. As a control, we used lentiviruses pseudotyped with a VSV glycoprotein, where no entry inhibition above 20% was detected for any of the drugs tested (data not shown). In sharp contrast, SARS-CoV-2 pseudoviruses were effectively blocked by most of the drugs previously tested on Vero E6 with wild-type virus (**Fig. 1D**). The main differences were observed with CA-074-Me, ciclesonide and arbidol. These compounds showed a partial blocking effect on ACE2 HEK-293T cells that was not obvious when using replication competent SARS-CoV-2 on Vero E6 (**Supp. Fig. 1**). In addition, NB-DNJ failed to block viral entry (**Fig. 1D**). Overall, using alternative SARS-CoV-2 viral systems, we could identify chloroquine derivatives, cathepsin inhibitors and cholesterol depleting agents as the most promising candidates to block SARS-CoV-2 endocytosis in Vero E6 and HEK-293T cells transfected with ACE2. However, chloroquine derivatives were the only ones that displayed an IC_50_ below 25 μM (**Table 1**), and were also active abrogating pseudoviral entry into HEK-293T cells expressing ACE2 (**Fig. 1E**). Although camostat failed to inhibit viral fusion on ACE2 HEK-293T cells (**Fig. 1E**), its activity was rescued when these cells were transfected with TMPRSS2. The opposite effect was observed for chloroquine, which reduced its inhibitory activity on TMPRSS2 transfected cells (**Fig. 1E**). Our results highlight that alternative routes govern SARS-CoV-2 viral entry and these pathways vary depending on the cellular target. Thus, effective treatments may need to block both plasma membrane fusion and endosomal routes to fully achieve viral suppression.

**Table 1.**
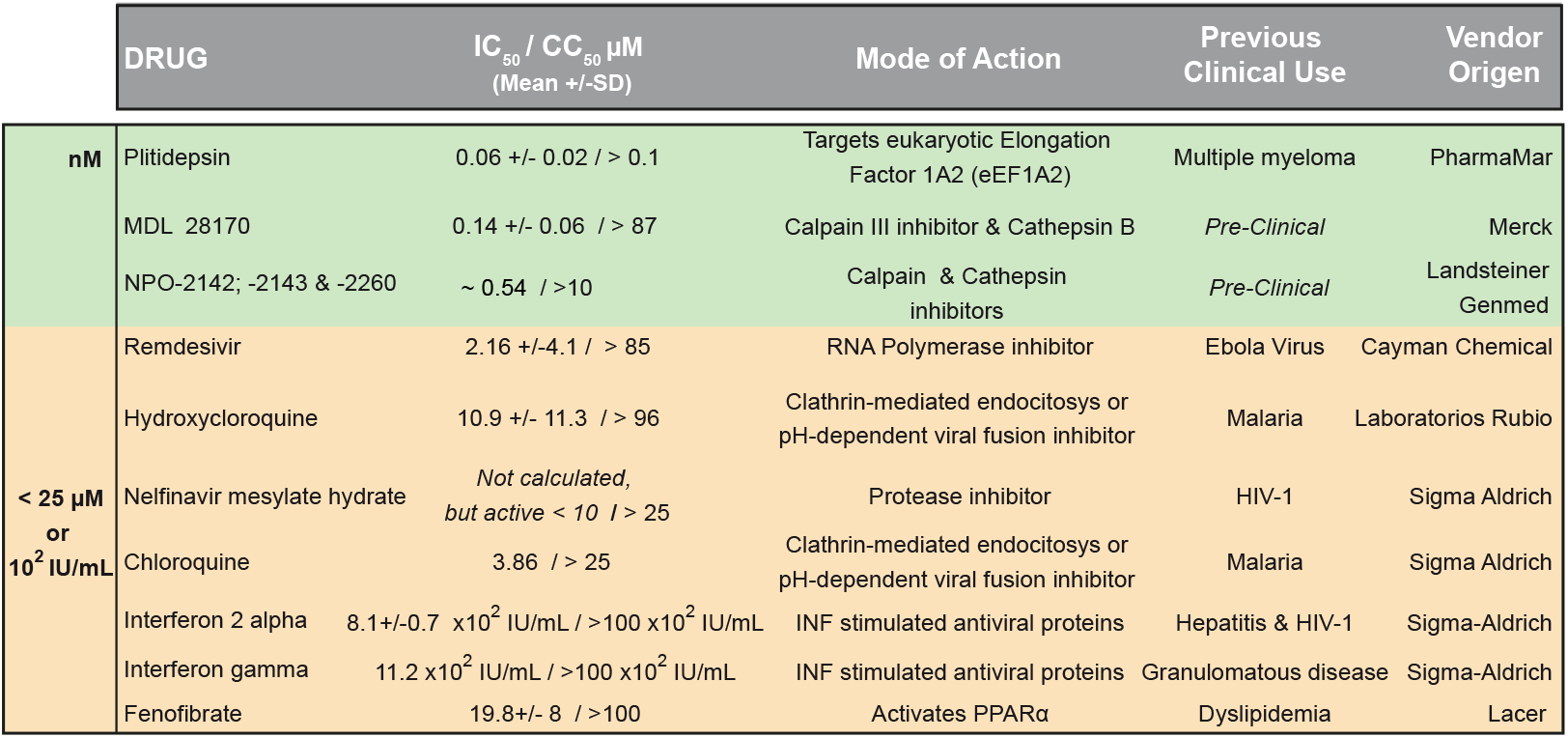
Compounds with antiviral activity grouped in colors depending on their IC_50_ values on Vero E6. Green color highlights compounds with the lower IC_50_ in the nM range, while orange color shows compounds in the μM or 10^2^ IU/mL range.

### 2. Antiviral activity of compounds that potentially inhibit post-entry steps

In our search for antivirals inhibiting post-viral entry steps, we first focused on remdesivir, which has *in vitro* activity against SARS-CoV-2 after viral entry^17^ and has already been approved for the treatment of COVID-19 by the FDA and EMA. We further confirmed the *in vitro* capacity of remdesivir to inhibit SARS-CoV-2-induced cytopathic effect on Vero E6 (**Fig. 2A**). The IC_50_ value of this drug in repeated experiments was always below 10 μM (**Supp. Table 2**). In combination with increasing concentrations of hydroxychloroquine, however, remdesivir did not significantly modified its own antiviral effect (**Fig. 2B**). This was also the case for other antivirals tested in combination (**Supp. Table 2**). Of note, other RNA polymerase inhibitors such as galdesivir, which was proposed to tightly bind to SARS-CoV-2 RNA-dependent RNA polymerase^38^, showed no antiviral effect (**Supp. Table 3**). Favipiravir, approved by the National Medical Products Administration of China as the first anti-COVID-19 drug in China^39^, showed only partial inhibitory activity at the non-toxic concentration of 100 μM (**Supp. Table 3**).

**Figure 2.**
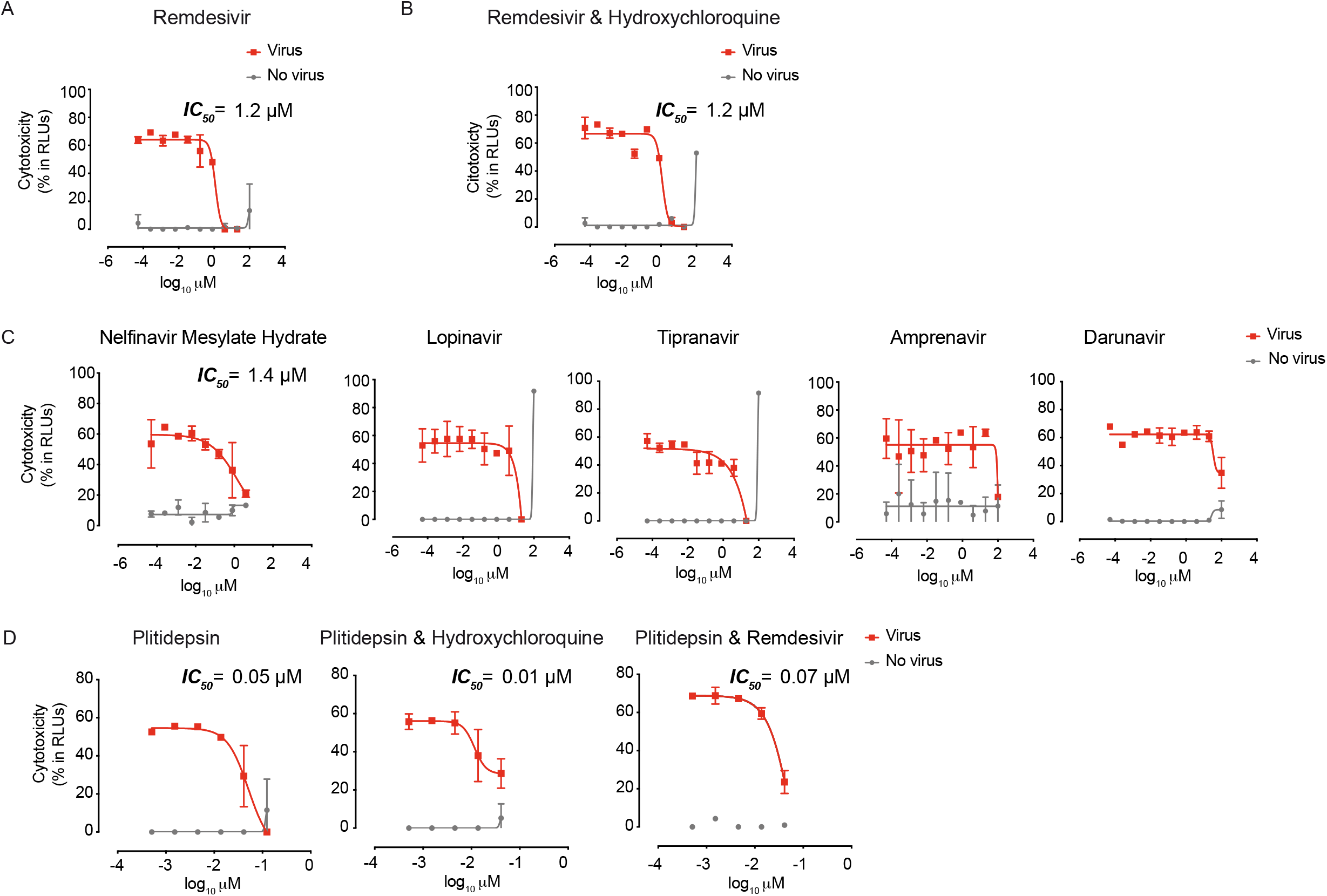
Antiviral activity of post-entry inhibitors. **A.** Cytopathic effect on Vero E6 cells exposed to a fixed concentration of SARS-CoV-2 in the presence of increasing concentrations of Remdesivir. Drug was used at a concentration ranging from 0.0512 nM to 100 μM. Non-linear fit to a variable response curve from one representative experiment with two replicates is shown (red lines), excluding data from drug concentrations with associated toxicity. The particular IC_50_ value of this graph is indicated. Cytotoxic effect on Vero E6 cells exposed to increasing concentrations of drugs in the absence of virus is also shown (grey lines). **B.** Cytopathic effect on Vero E6 cells exposed to a fixed concentration of SARS-CoV-2 in the presence of increasing concentrations of remdesivir and its combination with hydroxychloroquine, as detailed in **A**. Drugs in combination were used at a concentration ranging from 0.0512 nM to 100 μM (left panel). **C.** Cytopathic effect on Vero E6 cells exposed to a fixed concentration of SARS-CoV-2 in the presence of increasing concentrations of protease inhibitors against HIV-1. Nelfinavir mesylate hydrate was the only drug with activity. Inhibitors were used at a concentration ranging from 0.0512 nM to 100 μM. The particular IC_50_ value of this graph is indicated **D.** Cytopathic effect on Vero E6 cells exposed to a fixed concentration of SARS-CoV-2 in the presence of increasing concentrations of plitidepsin and its combinations with hydroxychloroquine and remdesivir. When combined, each drug was added at the same concentration. Drugs were used at a concentration ranging from 0.5 nM to 10 μM. The particular IC_50_ value of these graphs is indicated.

We also assessed clinically approved protease inhibitors with potent activity against HIV-1. However, none of the HIV-1 protease inhibitors detailed in **Supp. Table 3** showed remarkable protective antiviral activity against SARS-CoV-2 infection on Vero E6 cells, with the exception of nelfinavir mesylate hydrate, which showed an IC_50_ value below 10 μM (**Supp. Table 3** and **Fig. 2C**). Lopinavir and tipranavir inhibited SARS-CoV-2-induced cytopathic effect at the non-toxic concentration of 20 μM, and amprenavir exhibited activity at the non-toxic concentration of 100 μM (**Fig. 2C**). Darunavir, which has been tested in clinical trials, showed partial inhibitory activity at 100 μM, although this concentration had 8.5 ± 6.2 % of cytotoxicity associated (**Fig. 2C**). Of note, we tested HIV-1 reverse transcriptase inhibitors such as tenofovir disoproxil fumarate, emtricitabin, tenofovir alafenamide, and their combinations, but they also failed to show any antiviral effect against SARS-CoV-2 (**Supp. Fig. 1**). These results indicate that future clinical trials should contemplate the limited antiviral effect displayed by these anti-HIV-1 inhibitors against SARS-CoV-2 *in vitro.*

We also assessed the inhibitory capacity of HCV inhibitors, but none showed any antiviral activity (**Supp. Table 3**). Of note, exogenous interferons 2 alpha and gamma displayed antiviral activity against SARS-CoV-2 (**Supp. Table 3**). In light of these results, we tested the inhibitory effect of the TLR 7 agonist vesatolimod that triggers interferon production. Although this agonist was not able to protect from the viral-induced cytopathic effect on Vero E6 (**Supp. Table 3**), as expected since it is an interferon-producer deficient cell line^40^, it could still be useful in other competent cellular targets. Since severe COVID-19 patients display impaired interferon responses^41^, these strategies may be valuable to avoid disease complication. In addition, we also assessed several compounds with the best computational docking scores among approved drugs against the 3CL protease of SARS-CoV-2, but none of them were effective to protect Vero E6 from viral induced cytopathic effect (**Supp. Table 4**).

The most potent antiviral tested was plitidepsin (**Fig. 2D**), which targets the eukaryotic Elongation Factor 1A2 (eEF1A2) and has been previously used for the treatment of multiple myeloma. The mean IC_50_ value of this drug in repeated experiments was always in nM concentrations (**Supp. Table 3**). In combination with other active antivirals, we did not observe a reduction on IC_50_ values (**Supp. Table 2**). This result indicates no significant synergy, but also highlights the possibility of using plitidepsin without reducing its antiviral activity in combined therapies (**Fig. 2D**), what could be relevant to avoid possible selection of resistant viruses. Overall, plitidepsin showed the lowest IC_50_ values of all the compounds tested in this *in vitro* screening (**Table 1**).

### 3. Antiviral activity of compounds with unknown mechanism of action

We also assessed the inhibitory capacity of several inhibitors and broad anti-bacterial, anti-parasitic, anti-malarial, anti-influenza and anti-fungal compounds, along with other pharmacological agents previously suggested to interfere with SARS-CoV-2 infection (**Supp. Table 5**). Such was the case of ivermectin, an FDA-approved broad spectrum anti-parasitic agent previously reported to inhibit the replication of SARS-CoV-2 *in vitro* as measured by RNA accumulation^42^

However, among these potential antivirals, only three types of molecules exerted detectable antiviral activity in our assay: itraconazole, fenofibrate, and calpain and cathepsin inhibitors such as MDL 28170 and NPO compounds. Itraconazole, an antifungal that may interfere with internal SARS-CoV-2 budding within infected cells^43^, displayed an IC_50_ value of 80 μM (**Fig. 3A** and **Supp. Table 5**). Fenofibrate is clinically used to treat dyslipidemia via activation of PPARα, and also inhibited the cytopathic effect exerted by SARS-CoV-2 on Vero E6 at 20 μM (**Fig. 3B** and **Supp. Table 5**). As fenofibrate is a regulator of cellular lipid metabolism, we made use of the luciferase-based viral entry assay to try to elucidate its mode of action. When lentiviruses pseudotyped with the spike protein of SARS-CoV-2 were added to ACE2-expressing HEK-293T cells in the presence of fenofibrate, viral entry was abrogated (**Fig. 3C**). The most potent agent found was MDL 28170, a calpain III inhibitor in a pre-clinical stage of development that displayed activity in the nanomolar range (**Fig. 3D** and **Supp. Table 5**), as previously identified *in vitro*^44^ Moreover, three out of four different calpain and cathepsin inhibitors named NPO showed potent antiviral activity too (**Supp. Figure 2**). Of note, in combination with other active antivirals, we did not observe a reduction on IC_50_ values of MDL 28170 (**Supp. Table 2**).

**Figure 3.**
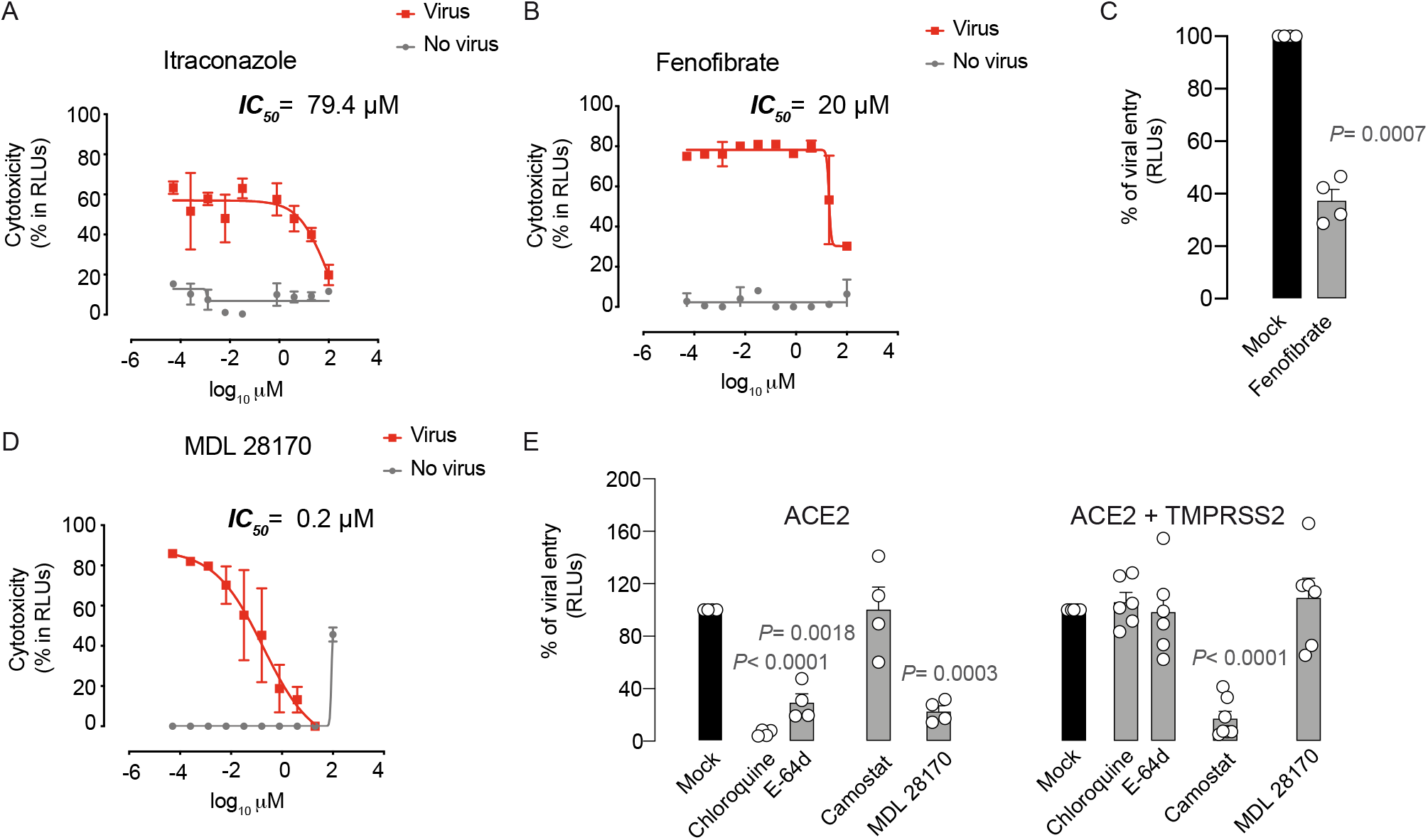
Antiviral activity of inhibitors with unknown mechanism of action. **A.** Cytopathic effect on Vero E6 cells exposed to a fixed concentration of SARS-CoV-2 in the presence of increasing concentrations of Itraconazole. Drug was used at a concentration ranging from 0.0512 nM to 100 μM. Non-linear fit to a variable response curve from one representative experiment with two replicates is shown (red lines), excluding data from drug concentrations with associated toxicity. The particular IC_50_ value of this graph is indicated. Cytotoxic effect on Vero E6 cells exposed to increasing concentrations of drugs in the absence of virus is also shown (grey lines). **B.** Cytopathic effect on Vero E6 cells exposed to a fixed concentration of SARS-CoV-2 in the presence of increasing concentrations of Fenofibrate, as detailed in **A. C**. Effect of fenofibrate on the entry of luciferase expressing lentiviruses pseudotyped with SARS-CoV-2 Spike in ACE2-expressing HEK-293T cells. Values are normalized to luciferase expression by mock-treated cells set at 100%. Mean and s.e.m. from two experiments with two replicates. Statistical deviations from 100% were assessed with a one sample t test. **D**. Cytopathic effect on Vero E6 cells exposed to a fixed concentration of SARS-CoV-2 in the presence of increasing concentrations of MDL 28170, as detailed in **A. E**. Comparison of MDL 28170 activity with entry inhibitors blocking viral endocytosis, such as chloroquine and E-64d, and inhibitors blocking serine protease TMPRSS2, such as camostat. ACE2 expressing HEK-293T cells transfected or not with TMPRSS2 were exposed to SARS-CoV-2 Spike lentiviruses in the presence of these compounds. Values are normalized to luciferase expression by mock-treated cells set at 100%. Mean and s.e.m. from at least two experiments with two replicates. Statistical deviations from 100% were assessed with a one sample t test.

Inhibitors of calpains, which are cysteine proteases, might impair the activity of viral proteases like 3CL (main protease) and PLpro (papain-like protease)^44,45^. However, calpain inhibitors may also inhibit cathepsin B-mediated processing of viral spike proteins or glycoproteins, including SARS-CoV and Ebola^45,46^. To understand the mechanisms of action of calpain and cathepsin inhibitors such as MDL 28170, we added lentiviruses pseudotyped with the spike protein of SARS-CoV-2 to ACE2-expressing HEK-293T cells and the same cells also expressing TMPRSS2 in the presence of this drug. Importantly, MDL 28170 only blocked viral entry in ACE2-expressing cells (**Fig. 3E**). This result indicates that MDL 28170 blocks cathepsins that are implicated in SARS-CoV-2 entry via the alternative endosomal pathway, as described for chloroquine derivatives and E-64d (**Fig. 3E**), which are all active when TMPRSS2 is not present and their inhibitor camostat displays no activity (**Fig. 3E**).

In conclusion, among the 72 compounds and their 28 combinations tested herein for their potential capacity to abrogate SARS-CoV-2 cytopathic effect, we only found 13 compounds with antiviral activity including camostat, and only eight types of these drugs had an IC_50_ below 25 μM or 10^2^ IU/mL (Table 1). These eight families of compounds were able to abrogate SARS-CoV-2 release to the supernatant in a dose dependent manner (Fig. 4), indicating that the reduction in the cytopathic effect that we had measured in cells correlates with viral production. As these eight families of compounds tackle different steps of the viral life cycle, they could be tested in combined therapies to abrogate the potential emergence of resistant viruses.

**Figure 4.**
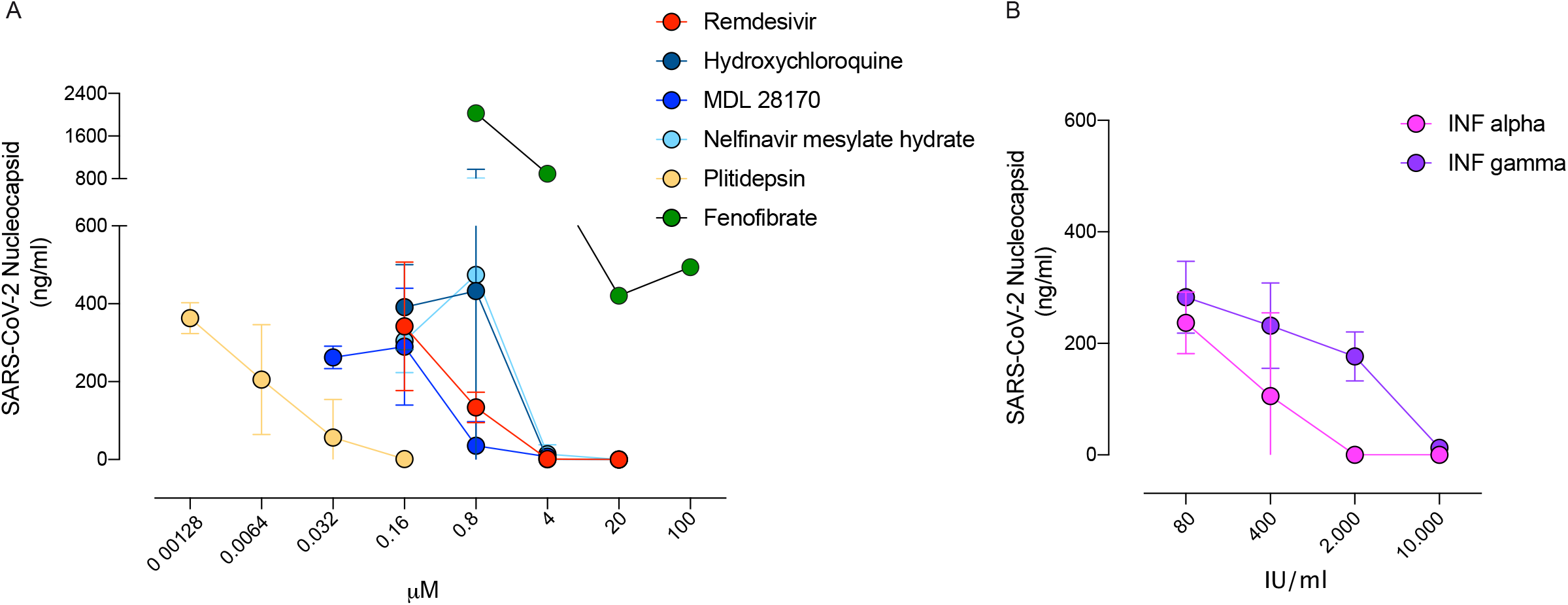
Decreased release of SARS-CoV-2 in the presence of inhibitors with antiviral activity. **A.** Viral release to the supernatant in the presence of the indicated compounds added at increasing concentrations 3 days post-infection of Vero E6 cells. SARS-CoV-2 nucleoprotein was detected with an ELISA at concentrations were drugs were nontoxic. Mean and s.e.m. from two experiments. **B.** Viral release to the supernatant in the presence of the indicated interferons as described in A. Mean and s.e.m. from one experiment.

## DISCUSSION

We have assessed the anti-SARS-CoV-2 activity of clinically approved compounds that may exert antiviral effect alone or in combination. Although we were not able to detect any remarkable synergy *in vitro,* combined therapies are key to tackle viral infections and to reduce the appearance of viral resistance. We have tested more than seventy compounds and their combinations, and verified a potent antiviral effect of hydroxychloroquine and remdesivir, along with plitidepsin, cathepsin and calpain inhibitors MDL 28170 and NPO, nelfinavir mesylate hydrate, interferon 2α, interferon-γ and fenofibrate. These are therefore the most promising agents found herein that were able to protect cells from viral-induced cytopathic effect by preventing viral replication.

Our findings highlight the utility of using hydroxychloroquine and MDL 28170 or other cathepsin inhibitors to block viral entry via the endosomal pathway in kidney cell lines such as Vero E6 or HEK-293T. However, the endosomal viral entry route is absent in pulmonary cells^18^ and, therefore, camostat should be considered as the primary inhibitor to limit SARS-CoV-2 entry in pulmonary tissues or in cells expressing TMPRSS2^6^. These findings can explain why randomized clinical trials using hydroxychloroquine have failed to show a significant protective effect^20,21^. Nonetheless, in combined therapies, it should be noted that agents targeting the alternative endosomal SARS-CoV-2 entry route such as hydroxychloroquine or MDL 28170 could be key to stop viral dissemination in other extrapulmonary tissues where viral replication has been already detected^47^, and viral entry could take place through this endosomal pathway. This could partially explain why in a retrospective observational study including more than 2500 patients, hydroxychloroquine treatment showed a significant reduction of inhospital mortality^48^. Thus, since alternative routes govern SARS-CoV-2 viral entry depending on the cellular target^49^, effective treatments might be needed to block both plasma membrane fusion and endosomal entry to broadly achieve viral suppression.

SARS-CoV-2 replication could be effectively blocked using nelfinavir mesylate hydrate, remdesivir and plitidepsin. While nelfinavir showed lower potency, remdesivir and plitidepsin were the most potent agents identified. However, remdesivir and plitidepsin are not yet suitable for oral delivery and require intravenous injection, complicating their clinical use for prophylaxis. Finally, we also confirmed the antiviral effect of type I and II interferons as well as fenofibrate, which have been extensively used in the clinic for many years and may therefore prove valuable for therapeutic use.

The data presented herein should be interpreted with caution, as the IC_50_ values of drugs obtained *in vitro* may not reflect what could happen *in vivo* upon SARS-CoV-2 infection. The best antiviral compounds found in the present study need to be tested in adequate animal models. This strategy already helped to confirm the activity of remdesivir against SARS-CoV-2^50^, while also questioning the use of hydroxychloroquine in monotherapy^18^. Thus, assessing antiviral activity and safety in animal models is key to identify and advance those compounds with the highest potential to succeed in upcoming clinical trials. In turn, *in vitro* results confirmed in animal models will provide a rational basis to perform future clinical trials not only for treatment of SARS-CoV-2-infected individuals, but also for pre-exposure prophylaxis strategies that could avoid novel infections. Prophylaxis could be envisioned at a population level or to protect the most vulnerable groups, and should be implemented until an effective vaccine is developed. In particular, orally available compounds with proven safety profiles, such as fenofibrate, could represent promising agents.

## ACKNOWLEDGEMENTS

We are grateful to patients at the Hospital Germans Trias i Pujol that donated their samples for research. For his excellent assistance and advice, we thank Jordi Puig from Fundació Lluita contra la SIDA. We are most grateful to Lidia Ruiz and the Clinical Sample Management Team of IrsiCaixa for their outstanding sample processing and management, and to M. Pilar Armengol and the translational genomics platform team at the Institut de Recerca Germans Trias i Pujol. We truly thank B. Trinité for generating the Spike expression plasmid construct used in this study. We thank Pharma Mar, Rubió Laboratories, Janssen and Drs. Cabrera and Ballana form IrsiCaixa; Pascual-Figal, Lax and Asensio-Lopez MC from “Instituto de Investigación Biosanitaria IMIB-Arrixaca” of Murcia; and Fernández-Real and Barretina from the “Institut d’Investigació Biomèdica de Girona Dr. Josep Trueta” for providing some of the reagents tested.

## FINANCIAL SUPPORT

The research of CBIG consortium (constituted by IRTA-CReSA, BSC, & IrsiCaixa) is supported by Grifols pharmaceutical. The authors also acknowledge the crowdfunding initiative #Yomecorono (https://www.yomecorono.com). JS, JVA and NIU have non-restrictive funding from Pharma Mar to study the antiviral effect of plitidepsin. The funders had no role in study design, data collection and analysis, decision to publish, or preparation of the manuscript.

## COMPETING INTEREST

A patent application based on this work has been filed (EP20382821.5). The authors declare that no other competing financial interests exist.

## SUPPLEMENTARY TABLES

**Supp. Table 1.** Antiviral activity of potential entry inhibitors tested against SARS-CoV-2. IC_50_ values are reported in μM unless otherwise indicated. “Not active” refers to the lack of inhibitory activity at the highest concentration tested for each compound.

| ACTIVITY | DRUG | IC<sub>50</sub> / CC<sub>50</sub> μM<br>(Mean +/-SD) | Mode of Action | Previous Clinical Use | Vendor Origen |
| --- | --- | --- | --- | --- | --- |
| ENTRY | Hydroxychloroquine | 9.3 +/- 11.1 / > 80 | Clathrin-mediated endocytosis or pH-dependent viral fusion inhibitor | Malaria | Laboratorios Rubió |
|  | Chloroquine | 3.9 / > 25 |  |  | Sigma Aldrich |
|  | Amantadine | *Not calculated, but partially active at 100 / > 100* | Clathrin-mediated endocytosis inhibitor | Parkinson & influenza A | Sigma Aldrich |
|  | Chlorpromazine (Largactil) | Not Active / > 18 | Clathrin-mediated endocytosis inhibitor | Antipsychotic | Sanofi |
|  | CA-074-Me | Not Active / > 34 | Cathepsin inhibitor B | *Pre-Clinical* | Sigma Aldrich |
|  | E-64d | *Not calculated, but partially active at 100 / > 100* | Cathepsin inhibitor B/L | *Pre-Clinical* | Sigma Aldrich |
| | Methyl-β-cyclodextrin | *Not calculated, but active at 1000* | Cholesterol-removing agent, lipid raft disruption | *Not approved* | Sigma Aldrich |
| | NB-DNJ | *Not calculated, but active at 100 / > 100* | Inhibits ceramide- glucosyltransferase and β-glucosidase 2 | Gaucher disease & Juvenile Sandhoff disease | Calbiochem |
|  | 3' Sialyllactose Na Salt | Not Active / 20 mM | Inhibits viral binding | *Pre-Clinical* | Carbosynth |
|  | Niclosamide | Not Active / > 9 | Beclin-1 stabilizer in autophagy | Helmints | Selleckchem |
|  | Ciclesonide | Not Active / > 20 | Glucocorticoid | Asthma | Selleckchem |
|  | Arbidol HCl | Not Active / > 40 | Fusion inhibitor? | Influenza | Selleckchem |
|  | Tofacitinib (Xeljanz) | Not Active / > 100 | JAK inhibitor | Rheumatoid arthritis | Pfizer |
|  | Baricitinib | Not Active / > 85 | JAK inhibitor | Rheumatoid arthritis | Selleckchem |
|  | Camostat | Not Active / > 100 | TMPRSS2 inhibitor | Chronic pancreatitis | Merck |
|  | Alpha-1 Antitrypsin | *Not calculated, but active at 12.5 mg/ml / > 12.5 mg/ml* | Cellular protease inhibitor | Alpha-1 antitrypsin deficiency | Grifols |

**Supp. Table 2.**
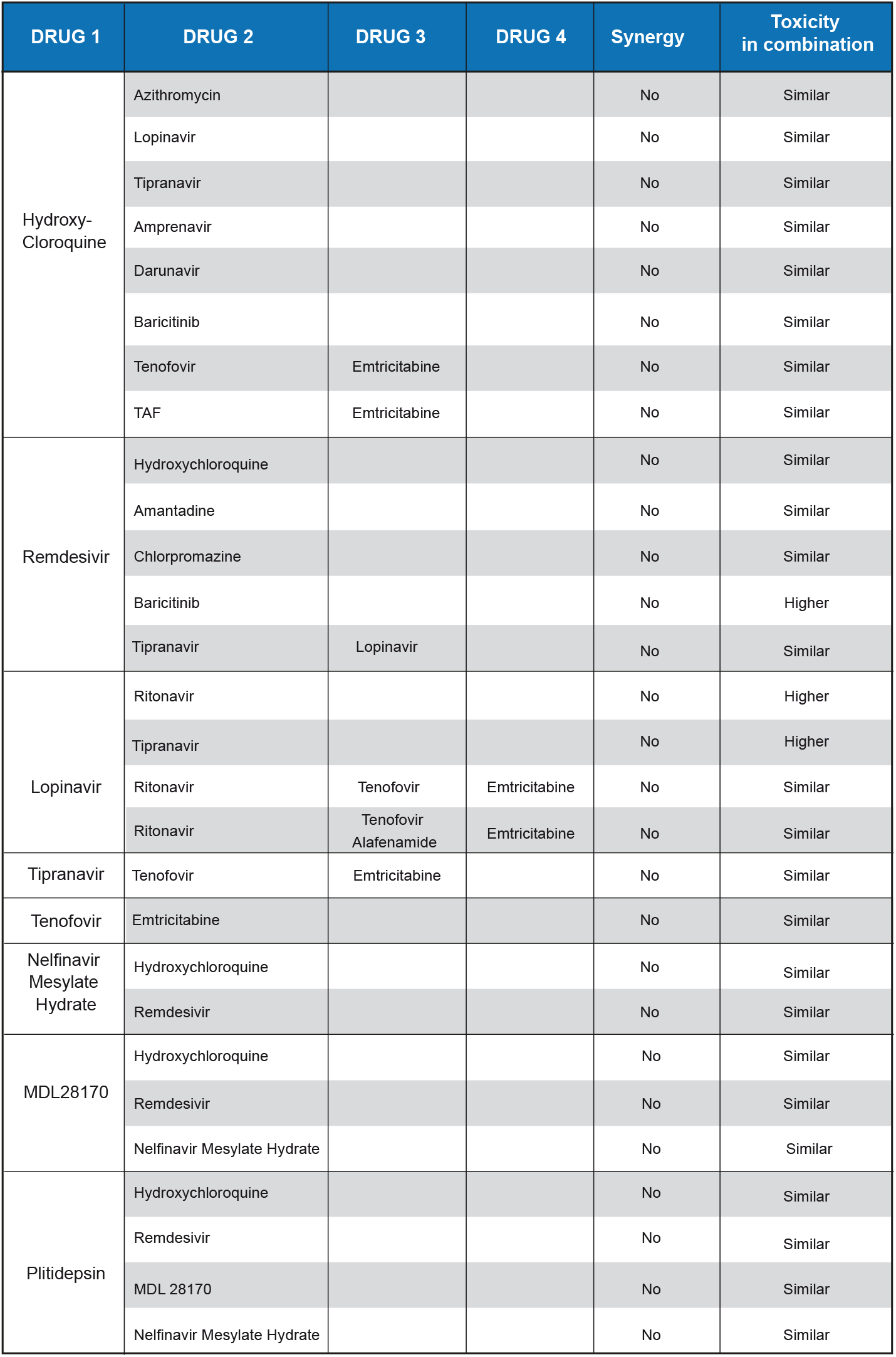
List of combinations of active antivirals that did not show any enhanced antiviral activity when mixed. Synergy was defined as showing an enhanced IC_50_ in drug combination as compared to the highest IC_50_ of the most potent compound tested. Similar toxicity is reported if viability was found to be similar to the most toxic compound tested in the combination, while higher toxicity reports an increase above 10% in cytotoxicity in the combination.

| DRUG 1 | DRUG 2 | DRUG 3 | DRUG 4 | Synergy | Toxicity in combination |
| --- | --- | --- | --- | --- | --- |
| Hydroxy-Cloroquine | Azithromycin |  |  | No | Similar |
|  | Lopinavir |  |  | No | Similar |
|  | Tipranavir |  |  | No | Similar |
|  | Amprenavir |  |  | No | Similar |
|  | Darunavir |  |  | No | Similar |
|  | Baricitinib |  |  | No | Similar |
|  | Tenofovir | Emtricitabine |  | No | Similar |
|  | TAF | Emtricitabine |  | No | Similar |
| Remdesivir | Hydroxychloroquine |  |  | No | Similar |
|  | Amantadine |  |  | No | Similar |
|  | Chlorpromazine |  |  | No | Similar |
|  | Baricitinib |  |  | No | Higher |
|  | Tipranavir | Lopinavir |  | No | Similar |
| Lopinavir | Ritonavir |  |  | No | Higher |
|  | Tipranavir |  |  | No | Higher |
|  | Ritonavir | Tenofovir | Emtricitabine | No | Similar |
|  | Ritonavir | Tenofovir Alafenamide | Emtricitabine | No | Similar |
| Tipranavir | Tenofovir | Emtricitabine |  | No | Similar |
| Tenofovir | Emtricitabine |  |  | No | Similar |
| Nelfinavir Mesylate Hydrate | Hydroxychloroquine |  |  | No | Similar |
|  | Remdesivir |  |  | No | Similar |
| MDL28170 | Hydroxychloroquine |  |  | No | Similar |
|  | Remdesivir |  |  | No | Similar |
|  | Nelfinavir Mesylate Hydrate |  |  | No | Similar |
| Plitidepsin | Hydroxychloroquine |  |  | No | Similar |
|  | Remdesivir |  |  | No | Similar |
|  | MDL 28170 |  |  | No | Similar |
|  | Nelfinavir Mesylate Hydrate |  |  | No | Similar |

**Supp. Table 3.** Antiviral activity of potential post-entry inhibitors against SARS-CoV-2. IC_50_ values are reported in μM unless otherwise indicated. “Not active” refers to the lack of inhibitory activity at the highest concentration tested for each compound.

| ACTIVITY | DRUG | IC<sub>50</sub> / CC<sub>50</sub> μM<br>(Mean +/-SD) | Mode of Action | Previous Clinical Use | Vendor Origen |
| --- | --- | --- | --- | --- | --- |
| POST-ENTRY | Remdesivir | 2.2 +/-4.1 / > 85 | RNA Polymerase inhibitor | Ebola Virus | Cayman Chemical |
|  | Galdesivir | Not Active / 100 | RNA Polymerase inhibitor | YFV | MedChemExpress |
|  | Favipiravir | *Not calculated, but partially active at 100 / > 100* | RNA polimerase inhibitor | Flavivirus, Arenavirus, Bunyavirus, Alphavirus | Quimigen |
|  | Saquinavir | Not Active / > 28 | Protease inhibitor | HIV-1 | Reference standard HPLC |
|  | Lopinavir | *Not calculated, but active at 20 / > 87* | Protease inhibitor | HIV-1 | Abbot |
|  | Ritonavir | *Not active / 20-100* | Protease inhibitor | HIV-1 | Abbot |
|  | Tipranavir | *Not calculated, but active at 20 / > 70* | Protease inhibitor | HIV-1 | Reference standard HPLC |
|  | Nelfinavir Mesylate Hydrate | *Not calculated, but active <10 / > 25* | Protease inhibitor | HIV-1 | Sigma Aldrich |
|  | Amprenavir | *Not calculated, but active at 100 / > 100* | Protease inhibitor | HIV-1 | GSK |
|  | Fosamprenavir Calcium | Not Active / > 25 | Protease inhibitor | HIV-1 | Sigma Aldrich |
|  | Darunavir | *Not calculated, but partially active at 100 / > 100* | Protease inhibitor | HIV-1 | Sigma Aldrich |
|  | Atazanavir Sulfate | Not Active / > 20 | Protease inhibitor | HIV-1 | Reference standard HPLC |
|  | Tenofovir disoproxil fumarate | Not Active / > 100 | Reverse Transcriptase inhibitor | HIV-1 | Selleckchem |
|  | Emtricitabine (Emtriva) | Not Active / > 100 | Reverse Transcriptase inhibitor | HIV-1 | Gilead |
|  | Tenofovir Alafenamide | Not Active / > 100 | Reverse Transcriptase inhibitor | HIV-1 | Selleckchem |
|  | Velpatasvir | Not Active / > 10 | linhibitor of NS5A protein | HCV | Selleckchem |
|  | Sofosbuvir | Not Active / > 100 | Polymerase inhibitor | HCV | Selleckchem |
|  | Boceprevir | Not Active / > 100 | Protease inhibitor | HCV | Quimigen |
|  | Vesatolimod | Not Active / > 20 | TLR7 agonist | Hepatitis & HIV-1 | MedChemExpress |
|  | Interferon 2 alpha | 8.1+/-0.7 x10<sup>2</sup> IU/mL / >100 x10<sup>2</sup> IU/mL | IFN stimulated antiviral proteins | Hepatitis & HIV-1 | Sigma-Aldrich |
|  | Interferon gamma | 11.2 x10<sup>2</sup> IU/mL / >100 x10<sup>2</sup> IU/mL | IFN stimulated antiviral proteins | Granulomatous disease | Sigma-Aldrich |
|  | Plitidepsin | 0.06 +/- 0.02 / > 0.1 | Targets eukaryotic Elongation Factor 1A2 (eEF1A2) | Multiple myeloma | PharmaMar |

**Supp. Table 4.**
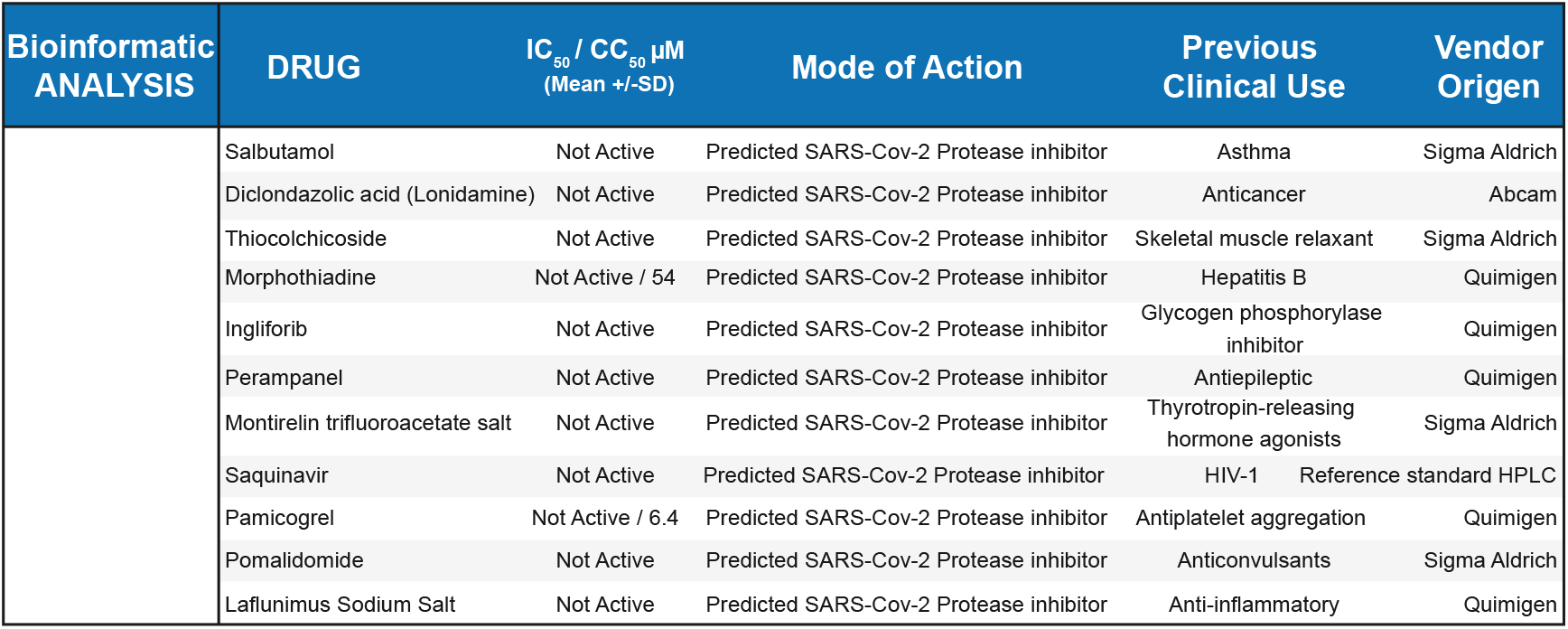
Antiviral activity of potential inhibitors against SARS-CoV-2 with predicted capacity to block SARS-CoV-2 viral protease. “Not active” refers to the lack of inhibitory activity at the highest concentration tested for each compound. All compounds but the two indicated were found non-toxic at 100 μM.

**Supp. Table 5.**
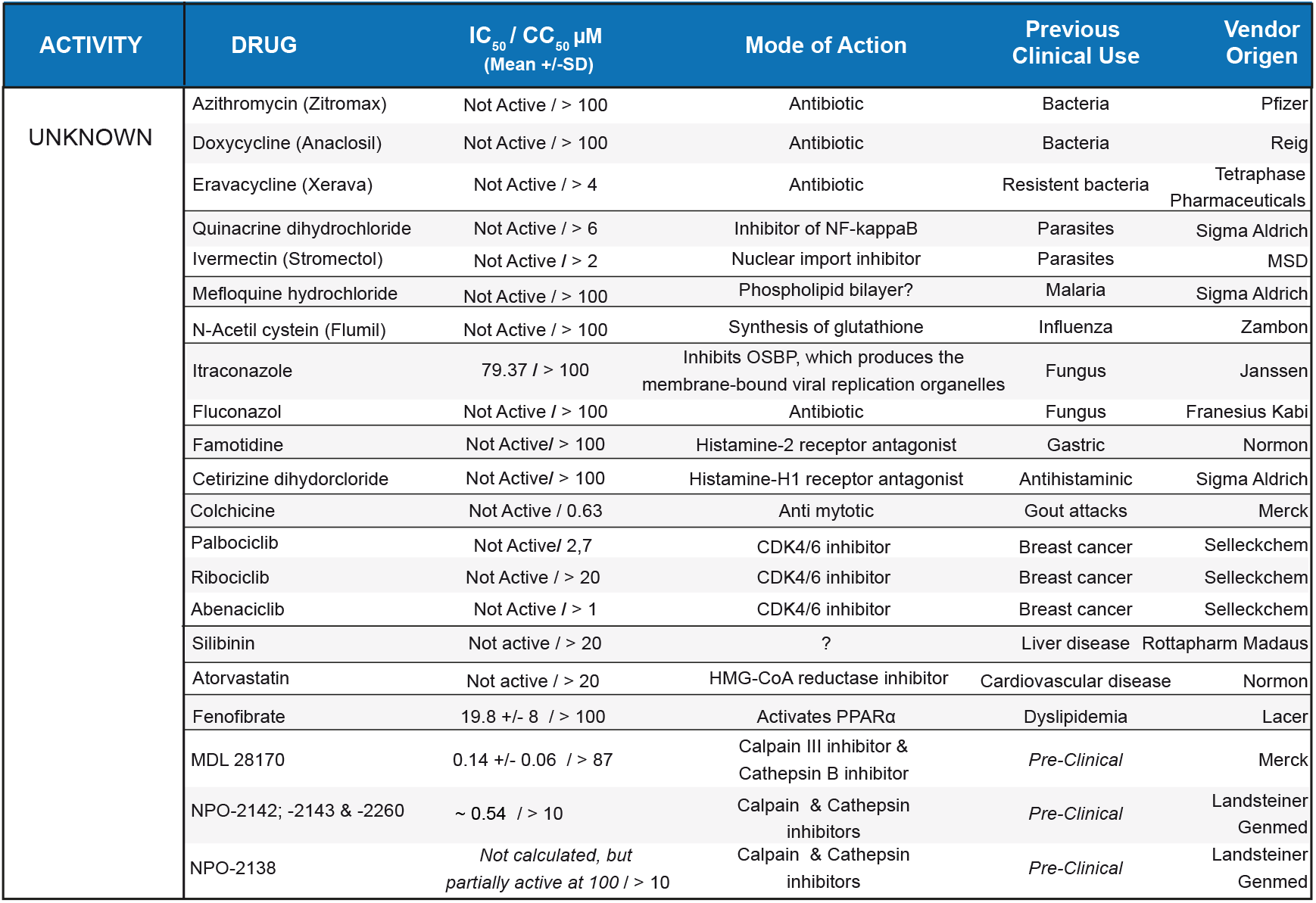
Antiviral activity of potential inhibitors against SARS-CoV-2 with unknown mechanism of action. IC_50_ values are reported in μM. “Not active” refers to the lack of inhibitory activity at the highest concentration tested for each compound.

## SUPPLEMENTARY FIGURES

**Supplementary Figure 1.**
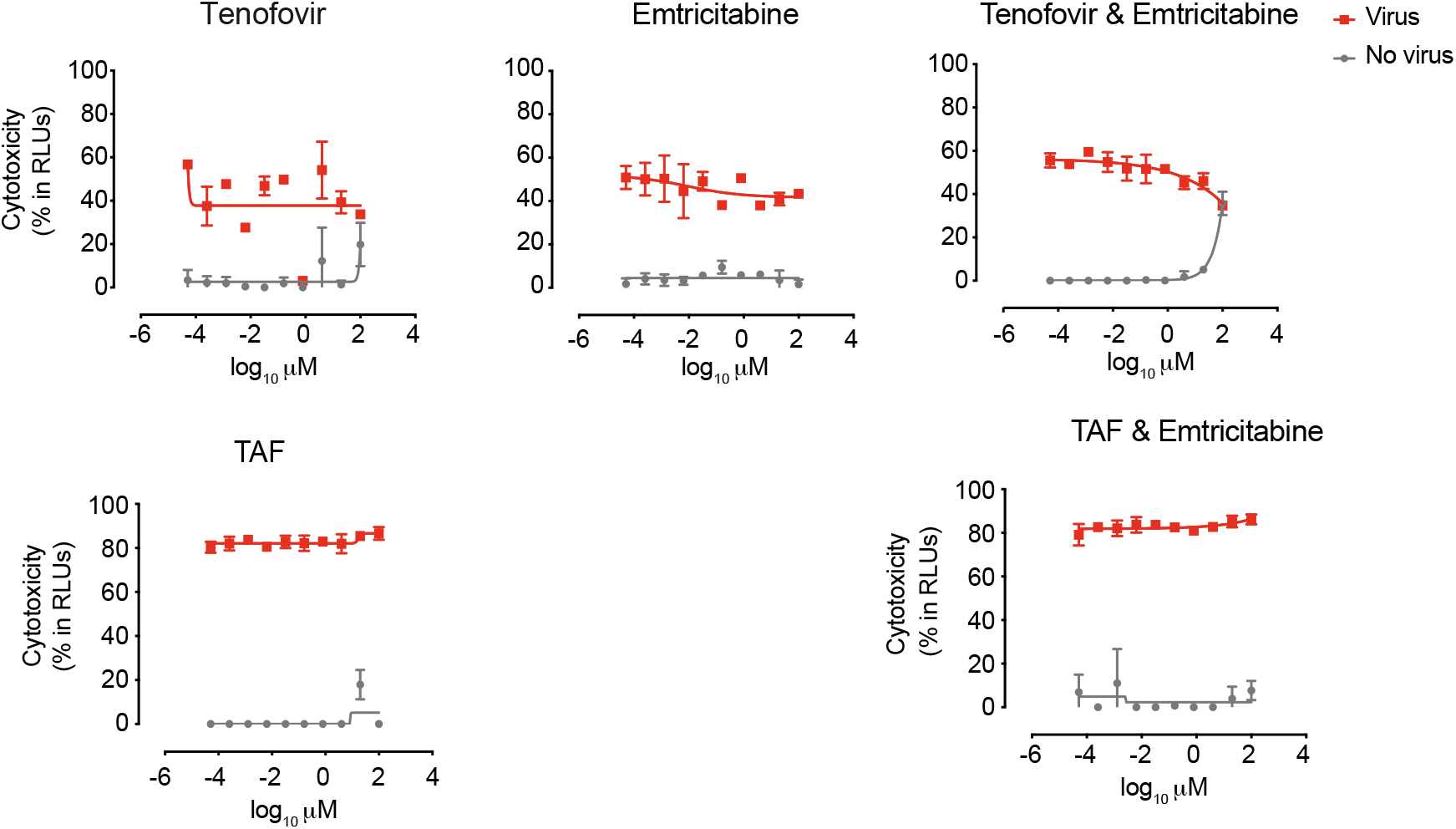
No antiviral activity of HIV-1 reverse transcriptase inhibitors. Cytopathic effect on Vero E6 cells exposed to a fixed concentration of SARS-CoV-2 in the presence of increasing concentrations of HIV-1 reverse transcriptase inhibitors. Drugs were used at a concentration ranging from 0.0512 nM to 100 μM. Non-linear fit to a variable response curve from one experiment with two replicates is shown (red lines). Cytotoxic effect on Vero E6 cells exposed to increasing concentrations of drugs in the absence of virus is also shown (grey lines).

**Supplementary Figure 2.**
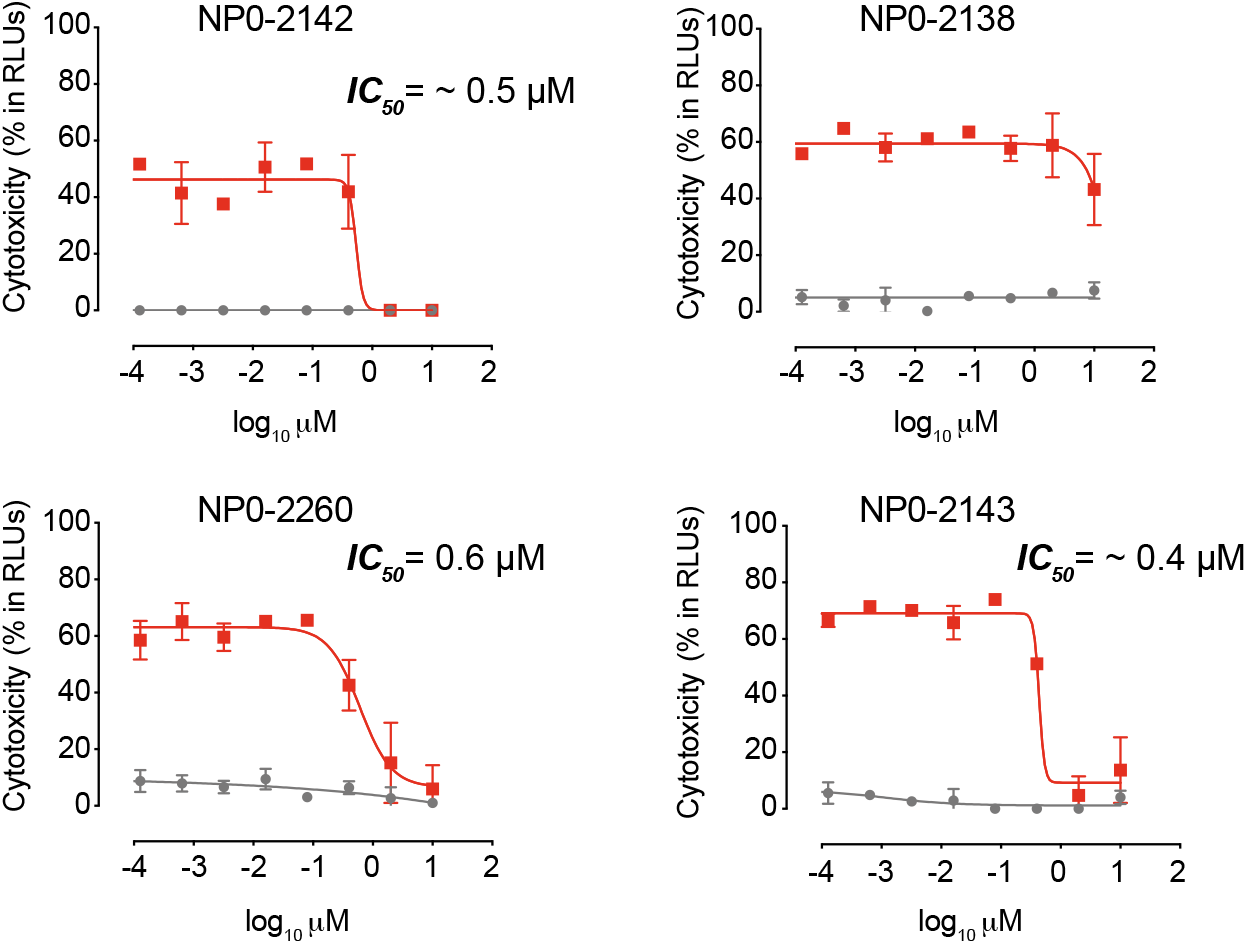
Antiviral activity of NPO calpain and cathepsin inhibitors. Cytopathic effect on Vero E6 cells exposed to a fixed concentration of SARS-CoV-2 in the presence of increasing concentrations of calpain and cathepsin inhibitors NPO. MDL 28170 was used at a concentration ranging from 0.0512 nM to 100 μM, while NPO was used from 0.00512 nM to 10 μM. Non-linear fit to a variable response curve from one experiment with two replicates is shown (red lines). Cytotoxic effect on Vero E6 cells exposed to increasing concentrations of drugs in the absence of virus is also shown (grey lines).

## AUTHOR CONTRIBUTION

Conceived and designed the experiments: JR, JMB, DPZ, JS, BC, JVA, NIU

Performed in silico drug modeling: VG

Performed experiments: JR, JMB, DPZ, MNJ, IE, VG, JVA, NIU

Contributed with critical reagents: CQ, IB

Analyzed and interpreted the data: JR, JMB, DPZ, MNJ, RP, LM, CQ, IE, IB, AV, VG,

JC, JB, JS, BC, JVA, NIU

Wrote the paper: JR, JVA, NIU

## DATA AVAILABILITY

Data is available from corresponding authors upon reasonable request.

## Notes

http://gisaid.org

